# Low intensity mechanical signals promote proliferation in a cell-specific manner: Tailoring a non-drug strategy to enhance biomanufacturing yields

**DOI:** 10.1101/2023.07.05.547864

**Authors:** M. Ete Chan, Lia Strait, Christopher Ashdown, Sishir Pasumarthy, Abdullah Hassan, Steven Crimarco, Chanpreet Singh, Vihitaben S Patel, Gabriel Pagnotti, Omor Khan, Gunes Uzer, Clinton T Rubin

## Abstract

Biomanufacturing relies on living cells to produce biotechnology-based therapeutics, tissue engineering constructs, vaccines, and a vast range of agricultural and industrial products. With the escalating demand for these bio-based products, any process that could improve yields and shorten outcome timelines by accelerating cell proliferation would have a significant impact across the discipline. While these goals are primarily achieved using *biological* or *chemical* strategies, harnessing cell mechanosensitivity represents a promising – albeit less studied – *physical* pathway to promote bioprocessing endpoints, yet identifying which mechanical parameters influence cell activities has remained elusive. We tested the hypothesis that mechanical signals, delivered non-invasively using low-intensity vibration (LIV; <1g, 10-500Hz), will enhance cell expansion, and determined that any unique signal configuration was not equally influential across a range of cell types. Varying frequency, intensity, duration, refractory period, and daily doses of LIV increased proliferation in CHO-adherent cells (+79% in 96h) using a particular set of LIV parameters (0.2g, 500Hz, 3x30 min/d, 2h refractory period), yet this same mechanical input *suppressed* proliferation in CHO-suspension cells (-13%). Exposing these same CHO-suspension cells to *distinct* LIV parameters (30Hz, 0.7g, 2x60 min/d, 2h refractory period) increased proliferation by 210%. Particle image velocimetry combined with finite element modeling showed high transmissibility of these signals across fluids (>90%), and LIV effectively scaled up to T75 flasks. Ultimately, when LIV is tailored to the target cell population, its highly efficient transmission across media represents a means to non-invasively augment biomanufacturing endpoints for both adherent and suspended cells, and holds immediate applications, ranging from small-scale, patient-specific personalized medicine to large-scale commercial bio-centric production challenges.

## Introduction

Biomanufacturing depends on the use of living cells cultured in bioreactors (Walsh, 2010).These cells are used to produce living biomaterials and therapeutic biomolecules, as well as to boost host-cell numbers *ex vivo* for personalized medicine. Current and future biomanufacturing applications include, but are not limited to, therapeutic protein production, treatment of cancer (Schuster et al., 2017), enhancement of the immune system, combatting infectious diseases, ameliorating metabolic dysfunction and building bioactive scaffolds for tissue engineering and regeneration (Lutolf & Hubbell, 2005; Nakashima & Reddi, 2003; Oberpenning et al., 1999; Reddi, 1998; Shea et al., 1999).

As demand for biomanufactured products increases, cost-effective optimization strategies are essential to improve yields in large-scale commercial production of therapeutic proteins, or more quickly expand cell numbers for personalized medicine applications including autologous immunotherapy. Bioreactor systems have become an indispensable element of this biomanufacturing process (Stephenson & Grayson, 2018), fostering a controlled biological, chemical and physical microenvironment to optimize cellular proliferation rate, stem cell differentiation, protein production and tissue development (Polak & Mantalaris, 2008). Thus, bioreactors are critical for providing not only a standardized, high-quality cell-based product, but for fostering a relevant yield of therapeutic cells (Eaker et al., 2016; Garcia-Aponte et al., 2021). However, the complexity of biological systems makes it difficult to design a generic bioreactor capable of controlling cell proliferation and functionality efficiently and across all cell types, a limitation that contributes to long biomanufacturing periods, disappointing yields and expensive therapies (Saini et al., 2021; Salehi-Nik et al., 2013).

Traditionally, optimization schemes using *biologic* strategies to improve cell yields involve invasive cell line development via vector genetic engineering, cell engineering or omics-based approaches to modulate and ultimately improve the transcriptional activities, the time integral of viable cell concentration and/or specific productivity (Gupta & Lee, 2007; Kim et al., 2012). From a *chemical* approach, culture media formulation and the chemical environment may be modulated, but jeopardize sterile culture systems by introducing or refreshing outside agents. The *physical* domain has also been shown to be important in bioprocessing, with a focus on substrate modulus (Özkale et al., 2021) and topology (Discher et al., 2005) or dynamic fluid motions (Banes et al., 1999), as a means of delivering mechanical cues to adherent cells (Ingber, 2008). Indeed, rocking, rotating and perfusion bioreactor systems that deliver fluid perturbations are commonly available and routinely used to maintain homogeneity across the media in favor of static cultures where cell expansion is comparatively slow. Despite the clear benefit of bioreactors, targeting specific cell types and biologic processes via tailored mechanical signals beyond a simple fluid agitation may require specialized technology, with the industry remaining unenthusiastic about processes that require expensive and time-consuming modifications to existing instrumentation and infrastructure (Kim et al., 2012; Stolfa et al., 2018). Therefore, there is a need to determine if one generic mechanical signal suits this purpose universally across distinct cell types, and if not, define cell-specific signals which promote rapid expansion during the culture phase of cell-based biomanufacturing (Garcia-Aponte et al., 2021; Li et al., 2010).

Low Intensity Vibration (LIV) is a dynamic mechanical signal characterized as low-magnitude (<1g peak-to-peak, where g = 9.8ms^-2^, Earth’s gravitational field) and delivered at a relatively high-frequency (10-500Hz). If shown effective, utilization of LIV promises a non-invasive, low cost and adaptable technology to augment proliferation rates in multiple cell types and could be readily integrated into existing bioreactor technology of varied design, particularly if the vibration signal transmits uniformly through the fluid. Herein, we explore LIV’s potential to promote cellular proliferation in both adherent and suspended cell culture systems. Beyond intensity and frequency, LIV variables include duration, dose number and refractory period.

LIV has been shown, *in vivo*, to protect bone quality and promote bone regeneration in mice (Rubin et al., 2007), rats (Rubin et al., 2001b), turkeys (Qin et al., 1998), sheep (Rubin et al., 2001a), and humans (Pagnotti et al., 2019). Applied to *in vitro* cell-based studies using adherent cells, LIV has been shown to influence lineage selection (Bas et al., 2020; Rubin et al., 2007), and promote expansion of mesenchymal stem cells (Chan et al., 2013; Frechette et al., 2017). Mechanistically, LIV acts through integrin signaling (Goelzer et al., 2020), including recruitment of focal adhesion kinase (FAK) and Akt to induce RhoA-mediated cell contractility (Uzer et al., 2015), as well as activating mechanically sensitive signaling molecules βcatenin (Uzer et al., 2018), and Yes1 Associated Transcriptional Regulator (YAP) (Goelzer et al., 2020; Thompson et al., 2020) that play interdependent roles in regulating cell proliferation in response to mechanical stimuli (Benham-Pyle et al., 2015; Sen et al., 2020; Zhao et al., 2008). Mounting evidence indicates that, as opposed to substrate strains and fluid shear stress that serve to distort and deform the cell membrane, LIV acts independent of substrate interaction by generating intracellular signals through the Linker of Nucleoskeleton and Cytoskeleton (LINC) complexes of the nuclear envelope by inertial forces generated by oscillatory accelerations (Birks & Uzer, 2021). While disabling LINC function is sufficient to mute LIV-induced signaling (Newberg et al., 2020; Touchstone et al., 2019; Xin Yi et al., 2020), neither LINC complex (Uzer et al., 2015), nor the nucleus itself (Graham et al., 2018) are necessary for activating the signaling events initiated by substrate strain. Consequently, LIV-induced effects on MSCs (Uzer et al., 2013), osteoblasts (Uzer et al., 2012) and osteocytes (Uzer et al., 2014) are largely independent of the LIV-induced fluid shear stress and substrate strain across multiple frequency/magnitude combinations.

Despite the demonstrated ability of LIV to influence adherent cell culture systems, it is unknown if the LIV signal, and the acceleration/deceleration of the cell, independent of substrate distortion, is applicable to cells grown in suspension. While many mechano-sensing pathways are conserved across cell types, it is entirely possible that suspension cells, such as Chinese hamster ovary (CHO) or T cells, would respond to LIV, or for that matter, require a mechanical signal distinct from adherent cells. CHO cells were chosen as our cell-type of interest, due to their prevalence in the pharmaceutical and biotech industries, and their ability to be cultured as either adherent or suspension cells (Baker et al., 1989). Our studies ultimately seek to determine if LIV represents a non-invasive, non-pharmacological engineering strategy to foster a proliferative response in cell types used in biomanufacturing without altering functionality or viability.

## Methods

### Delivery of LIV via a Feedback-Controlled System

Originally designed for *in vivo* use, low magnitude, high frequency, mechanical signals are delivered using LIV via a closed-loop acceleration feedback-controlled system (Rubin et al., 2004). For *in vitro* cell culture studies, the LIV system was modified to accommodate cell culture vessels (*e.g.,* microplates and T75 flasks), and be used under sterile cell culture conditions including a high humidity environment within an incubator (**Fig. 1**). The LIV device is controlled by an electromagnetic actuator, generating vertical oscillations, while a damped-spring/slider system ensures a smooth sinusoidal signal. The LIV signal is monitored with a platen-mounted accelerometer and controlled through current driving closed-loop error feedback proportional-integral-derivative (PID) controller (*LabVIEW NI*, TX).

**Fig. 1:**
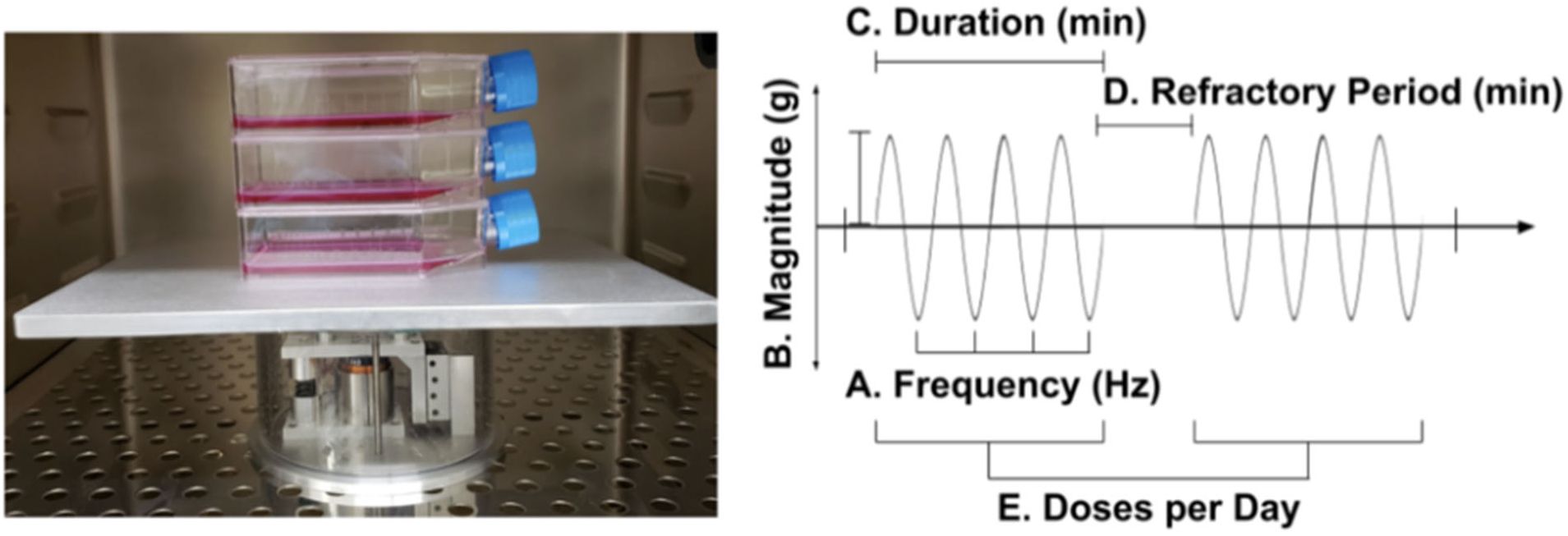
LIV device pictured inside incubator with stacked T75 flasks on the top platen (left). Driving the platform is an electromagnetic actuator displacing in the vertical direction, with current regulated through closed-loop feedback as based on an accelerometer affixed to the bottom of the top platen. LIV parameters examined in these protocols (right) are customizable via LabView PID, and include magnitude, frequency, duration, refractory period and doses per day.

To protect from humidity, all electronic components were housed in an airtight plexiglass enclosure, enabling 24/7 automated regulation and monitoring. LIV signals were delivered to cell culture vessels at prescribed magnitudes and frequencies depending on the signal required for each experiment, ranging from 0.1g-1.2g peak-to-peak and 10Hz-500Hz, respectively (± 5%). Starting parameters for frequency and magnitude for the LIV signals were selected based upon previous data from *in vitro* studies of other adherent cell types such as MSC (Uzer et al., 2016), pre-osteoblasts and osteocytes (X. Yi et al., 2020). Additional parameters included the number of LIV bouts per day, signal duration per bout, and refractory period (time elapsed between bouts; **Fig. 1**) (Sen, Xie, et al., 2011).

### Quantifying the Transmissibility of LIV signals to Suspended Cells

In order to determine LIV-induced fluid motions and transmissibility of the signal across culture media in 6-well plates, we used a finite element model (FEM) to derive LIV-induced fluid shear stresses *in vitro* during 0.7g, 90Hz vertical vibration (Abaqus 6.9.1, Simula, RI).(Uzer et al., 2012) The culture well was modeled as a deformable shell element (Polystyrene E=3GPa). Vertical vibrations were applied as a velocity boundary condition to cylindrical walls while the well bottom was not constrained and was free to deform. A Eulerian-Lagrangian contact algorithm was used to model fluid motion due to LIV. A conceptual schematic and the analysis region of interest are shown in **Figure 2**.

**Figure 2:**
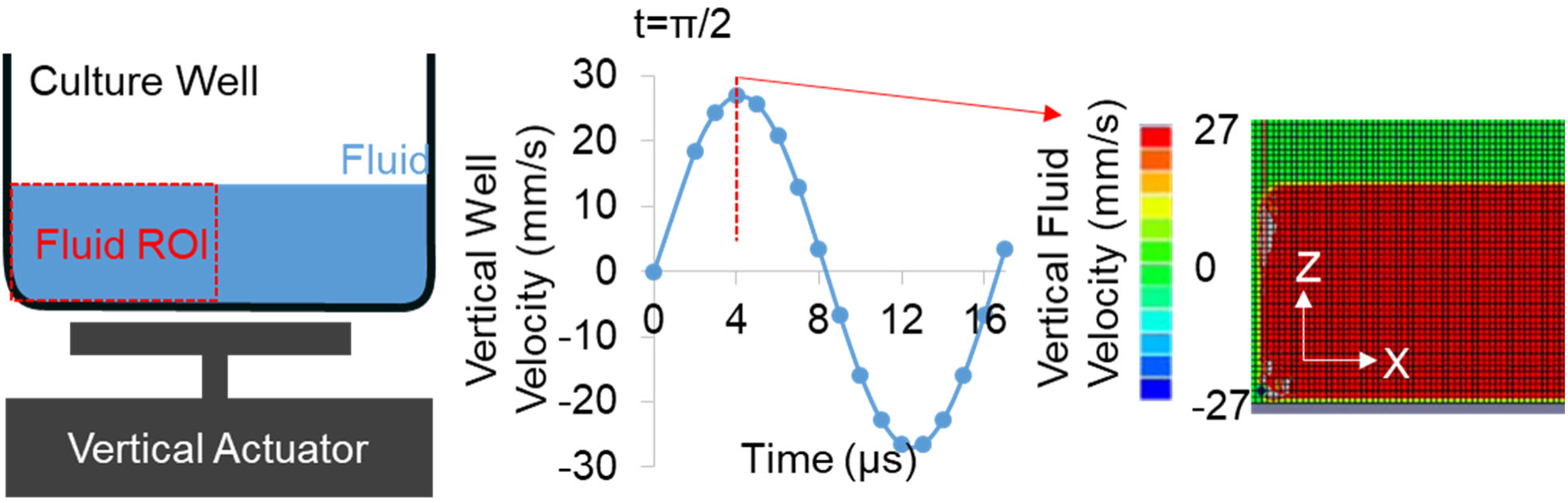
A finite element model was used to determine fluid motions in a vertical vibration of a 6-wellplate (left). Peak out of plane strain(ε) during 2 cycles of 0.7g, 90Hz vertical vibration is shown in center. Fluid motion was modeled in Abaqus (6.9.1, Simula, RI). Vertical velocity distribution during t =π/2 shows that fluid motion was coupled to the vertical well peaking at 27mm/s (right).

To explore the feasibility of scaling up the overall cell expansion using larger culture vessels, LIV signal transmissibility and fluid motion was also experimentally measured in vertically oscillating T75 vessels (**Fig. 3**), comparing completely filled flasks to manufacturer suggested partially filled fluid volumes using particle image velocimetry (Uzer et al., 2012).

**Figure 3.**
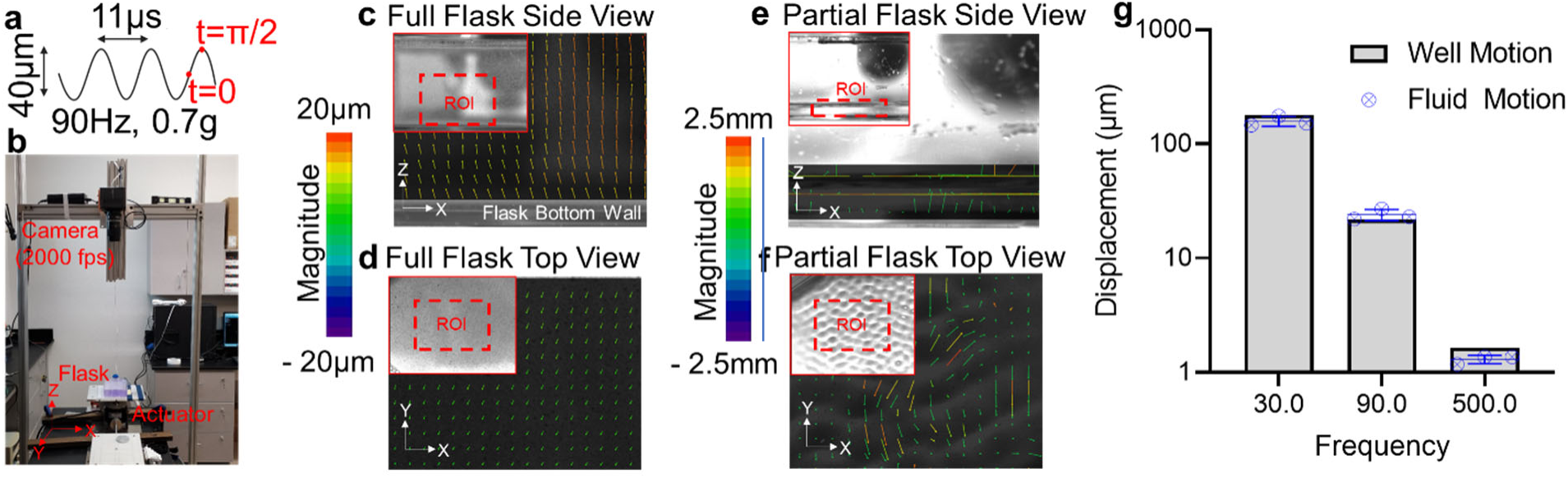
– Full or partially filled flasks were vibrated and fluid motions were quantified. (a) LIV was applied at 90Hz and 0.7g, generating peak motion of 20µm per cycle. (b) A Photron UX50 high speed camera at a rate of 2000 frames per second recorded the fluid motion from either side view or top view during vertical vibration driven by a Labworks ET-126HF-1,-4 (13lbf) transducer using a sinusoidal driving function. Frames between t=0 and t=π/2 that correspond to peak cycle displacement were compared within the region of interest (ROI, red bounding box) to determine the acceleration transmittance and fluid sloshing. (c) Side motion of the full flasks shown that fluid motion vectors were lined parallel to actuator motion with a maximum magnitude of 18µm, essentially matching to the 20µm peak motion of the actuator and (d) fluid sloshing measured from top view was minimal around 2µm. (e) While displacement vectors from the side view of partially filled flasks showed peak motion at 2.5mm – two orders of magnitude larger than filled flasks. (f) Similarly, fluid sloshing measured from top view showed large motions of unconstrained fluid motions peaking at magnitude of 2.5mm. (g) Comparing the transmittance of 0.7g acceleration magnitude in full flasks shows close to 100% transmittance across 30, 90 and 500 Hz frequencies.

Speckles with 1g/mL density (Cospheric, VIOPMS 63-75um) were suspended for tracking fluid motion. To track light-reflective speckles, a white LED light source was used, and the motion of the surface was captured with a high-speed camera (Photron UX50, San Diego, CA) at a rate of 2000 frames per second (fps). Culture vessels were vibrated vertically at 90Hz with an acceleration of 0.7g peak-to-peak, requiring a vertical displacement of approximately 20µm. Recording of sample motion was started 15s after the start of LIV to ensure that steady state was reached. Recorded high speed videos at 2000fps were analyzed using previously developed CASI software (Uzer et al., 2012). For this analysis we have compared the fluid motion differential between t=0 and t =π/2 time points that correspond to the peak cycle displacement of 20µm. This analysis was repeated for 30Hz and 500Hz using filled T75 flasks to experimentally quantify the transmissibility of LIV signals to cells cultured in suspension in larger volume conditions.

### CHO-Adherent Cell Culture

CHO-adherent cells were cultured in an incubator at 37°C and 5% CO_2_ (cells were provided by G. Balazsi at Stony Brook University). Prior to the experiment, on day 0, cells were plated in 6-well tissue culture treated plates at the concentration of 6x10^4^ cells/ml in 2ml of supplemented Ham’s F-12K (89% Ham’s F-12K, 10% FBS, 1% penicillin/streptomycin; *Life Technologies*, NY) per well. Cells were allowed to attach to the surface and grow overnight. Starting 24h after plating, cells were either subject to LIV (V; n=6) or sham-handled (NV; n=6) for 3d. On day 4, cells were trypsinized (0.05% Trypsin-EDTA; *Life Technologies, NY*) and counted with 1:1 ratio of cell solution: Trypan Blue (*Life Technologies*) to assess cell viability using an automated cell counter (Countess II FL; *Thermo Fisher Scientific*, MA).

### CHO-Suspension Cell Culture

CHO-suspension cells were plated at a concentration of 7.5x10^4 cells/mL in Freestyle CHO Expression medium supplemented with 8 mM L-glutamine (*Thermo Fisher Scientific*, MA) in 125-mL disposable polycarbonate Erlenmeyer flasks with vented caps (*Fisher Scientific,* NH) and cultured in a CO_2_ cell incubator at 37°C and 8% CO_2_. Starting 24h after plating, cells were either subjected to LIV (V; n=6) or sham-handled (NV; n=6) for 3d. Cells were counted with 1:1 ratio of cell solution: Trypan Blue (*Life Technologies*, NY) using an automated cell counter (Countess II FL; *Thermo Fisher Scientific*, MA).

### T Cell Culture

Human-derived CD3+ Pan T cells, CD4+ T cells, and CD8+ T cells (*HemaCare*, NY) were obtained from Caucasian male donors. CD3+ cells were isolated using negative selection, and all were cultured in 24-well plates or 6-well plates, with supplemented RPMI (95% RPMI, 5% FBS, 1% penicillin/streptomycin: *Life Technologies*, NY) at 37°C and 5% CO_2_. Seeding concentrations for all culture vessels were 0.5x10^6^ cells/mL with Dynabead^®^ CD3/CD28 (*Thermo Fisher Scientific*, MA) and recombinant interleukin-2 (rIL-2; 10ng/mL) for activation. During the initial activation phase, Dynabeads^®^ were at a 1:2 cell:bead ratio. Once a sufficient cell number had been reached to begin experimentation, Dynabeads^®^ were removed using a DynaMag^TM^ (*Thermo Fisher Scientific*, MA) magnet and replenished at a 1:10 cell:bead ratio 24h prior to the start of the experiment.

Following plating, T cells were either subjected to LIV (V) or sham-handled (NV). Cell counts were performed every 24h prior to receiving LIV doses on representative aliquots. Dynabeads^®^ were removed before mixing the cell suspension with Trypan Blue at a 1:1 ratio of cell solution:Trypan Blue. Counts were taken on each sample using an automated cell counter (Countess II FL; *Thermo Fisher Scientific*, MA). The average of these counts was considered the cell number for the day, and the milestone of proliferation.

### Statistical Analysis

Data normality was assessed with a Shapiro-Wilk test (α= 0.05). Depending on the normality of the data, either a Student’s t-test or Mann-Whitney U test were performed to measure the mean difference between two groups (*GraphPad Prism*, CA). A Student’s t-test was used in situations where the data were normally distributed, and the Mann Whitney U test in cases where the dataset was found to be non-normally distributed. In the case of analyzing the mean difference between at least three groups, datasets with a normal distribution were analyzed with a one-way ANOVA (Turkey’s post-hoc test), and non-normally distributed data were analyzed using the Kruskal-Wallis test (Dunn’s post-hoc test) (*GraphPad Prism,* CA). All tests were performed with α=0.05, a power of 95% with p<0.05 being considered statistically significant.

## Results

### Transmission of LIV Across Media to Suspended Cells

In 6-well plates, the peak surface strain at t = π/2 was 1.4με (microstrain). Comparing the peak velocities of both the 6-well wall and the fluid showed a matching velocity of 27mm/s (**Figure 3a**). Further, fluid velocity showed no large gradients across the well height indicating that the attached and suspended cells experience lossless LIV transmittance from the actuator. While the lossless transmission of LIV in 6-well plate is largely due to small plate deformations due to well-supported and small surface area (∼10cm^2^), T75 flasks have considerably larger surface (75 cm^2^) and result in larger effective deformations during LIV. As reported by our group (Uzer et al., 2015) and others (Cox et al., 2005; Dareing et al., 2007), bending motions in a vertically oscillating elastic culture plate will result in fluid motions that will propagate to the fluid surface thus altering the acceleratory motions experienced by suspended cells. A common strategy to limit fluid motions is to fill the culture vessel to minimize differential accelerations between gas and liquid mediums (Thompson et al., 2020). Therefore, to test whether filling a culture vessel maximizes the transmittance of vertical accelerations to fluid we quantified the acceleration transmittance in fluid-filled, vertically-oscillating culture vessels and compared it to manufacturer suggested fluid volumes (**Figure 3b**).

Transmittance was compared from the side view (X-Z plane) while the fluid motion at the surface was compared from the top view (X-Y plane). Side motion of the full flasks is shown in **Figure 3c**, fluid motion vectors were parallel to each other and to the actuator motion with a maximum magnitude of 18µm, essentially matching to the 20µm peak motion of the actuator, indicating that fluid particles were vibrating at the same frequency and acceleration magnitude as the actuator. Shown in **Figure 3d**, fluid sloshing measured from top view was minimal around 2µm, again indicating minimal fluid motion in filled flasks. Measuring fluid motions at the fluid/flask boundary in partially filled flasks, we have observed that fluid was forming standing wave patterns and showed vertical motions across the measured area (**Figure 3e**). Fluid displacements during half cycle (0.05s) reached up to 250µm, indicating fluid towards the surface was moving approximately 10 times more than fluid filled flasks. Similarly, fluid movement measured from top view showed motions peaking at approximately 2.5mm with the fluid surface in a standing wave pattern (**Figure 3f**), suggesting that mechanical signals generated by LIV are a product of both acceleration and fluid motion, and each need be considered when calculating acceleration transmittance. LIV Transmissibility of the LIV signal across fluid in filled flasks at 0.7g acceleration exceeded 95%, and was highly similar between signals operating at 30, 90 & 500Hz (**Figure 3g)**, suggesting that LIV-induced harmonic motions of the flask surface were directly transmitted to the fluid with minimal loss across wide ranges of frequencies.

### Proliferative LIV Signal for Adherent Cells Hampers Growth for suspension cells

To minimize potential differences between our representative adherent and suspension cells we compared CHO cells grown as adherent cells to those grown as suspension culture conditions. We first tested the effect of a LIV signal, first optimized for use with MSCs, on adherent CHO cells. Control and experimental cells were plated on Day 0 and allowed to adhere to the plate overnight. 24h after plating, adherent-CHO cells were subjected to a LIV signal for 3d that was delivered at 0.2g and 500Hz in 3x30min/d doses with a 2hr refractory period between each dose. At 4d, cells were counted, and cell numbers compared against a sham LIV control. Following 96h, CHO adherent cells subject to the LIV program showed a 79% ± 30.7% increase in cell number compared to sham controls (n=6, p<0.05; **Figure 4a**).

**Fig. 4:**
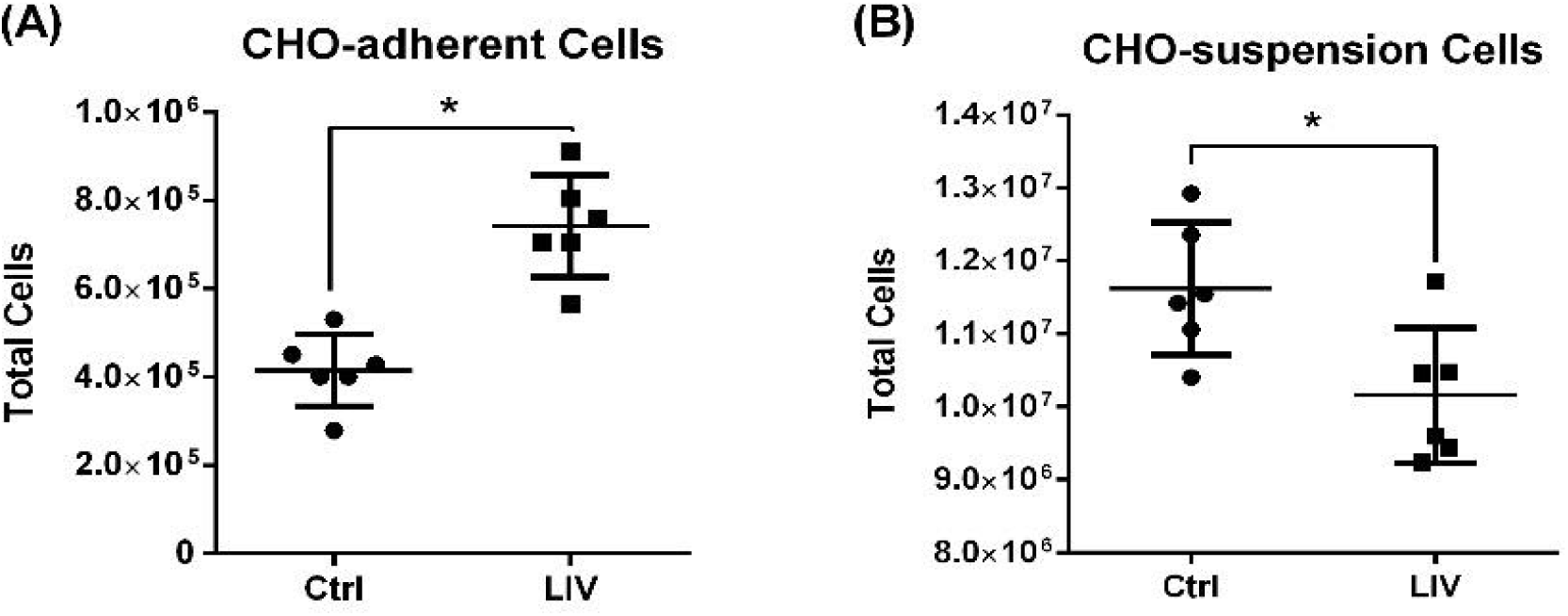
LIV signal of identical intensity, frequency, duration and refractory period (0.2g, 500 Hz, 3X30 min/d, 2h refractory period) showed significant, but opposing, proliferative responses in (A) CHO-adherent cells (+79%) and (B) CHO-suspension cells (-13%) (n=6 per group, mean ± SD, p≤0.05)

We delivered the same LIV signal parameters to CHO-suspension cells. Plated on Day 0 and allowed to grow overnight, LIV cells were then subjected to the identical signal as the CHO-adherent cells (0.2g, 500Hz, 3x30min/d doses, 2hr refractory period), and compared to sham LIV suspension controls. At 96h, LIV-treated CHO-suspension cell population had fallen -13% ± 10.1% below sham LIV controls (n=6, p<0.05; **Figure 4b**). While these data confirm that CHO-suspension cells are mechanosensitive, they also suggest that LIV signal parameters are cell-type specific, and that the same set of parameters that enhance proliferation in adherent cells hampers proliferation in suspension cells.

### LIV Signal Optimization for Suspension Cells

To determine whether a LIV signal could be designed to enhance proliferation of suspension cells, a series of experiments were conducted to examine both frequency and intensity of the LIV signal. To determine the optimal frequency for enhancing proliferation, 5 sets of CHO-suspension cells were plated and allowed to culture overnight. Each set of CHO-suspension cells was subjected to a LIV signal with a consistent amplitude of 0.2g, while the frequency was varied between 0Hz (sham-LIV control), 30Hz, 60Hz, 90Hz, and 250Hz between groups. Following 3d exposure to LIV, cell counts were taken from each group. The 30Hz frequency increased CHO-suspension cell number by 61% ± 10.1% relative to sham LIV control, while no other frequency showed significant difference to the control (n=6, p<0.05; **Figure 5a**).

**Fig. 5:**
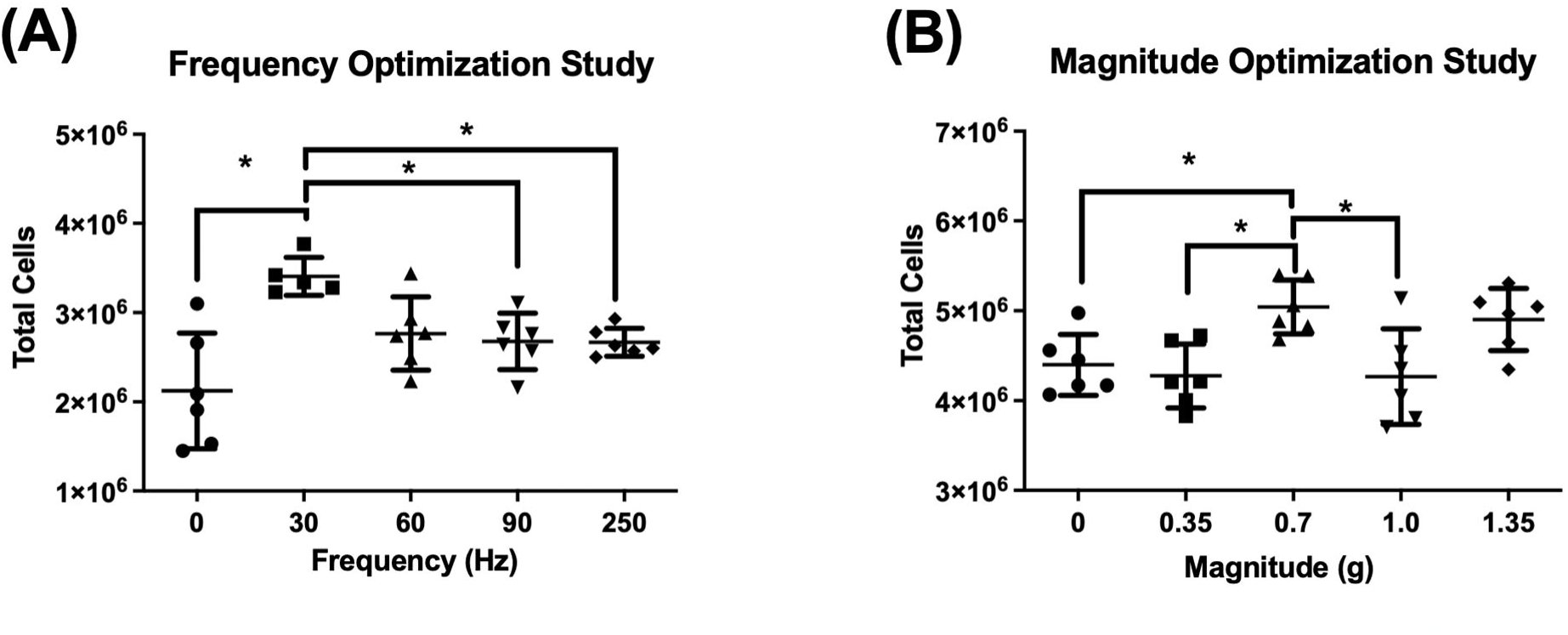
(A) Altering the LIV frequency applied to CHO-suspension cells showed a 61% increase at 30Hz compared to control, significantly greater than 90Hz or 250Hz. (B) Then, altering magnitude while at 30Hz, 0.7g was distinguished as more influential than other inputs with a 15% increase compared to control. (n=6 per group, mean ± SD, p≤0.05).

To determine which LIV intensity would promote the largest increase in CHO-suspension cell proliferation, 5 sets of CHO-suspension cells were plated and allowed to culture overnight. Each set of CHO-suspension cells was then subject to a LIV signal using the ‘optimized’ frequency of 30Hz, with acceleration adjusted to include 0g (sham-LIV control), 0.35g, 0.7g, 1.0g, and 1.35g. After 3d of LIV, cell counts were taken from each group. An acceleration of 0.7g generated the largest increase in proliferation of 15% ± 6.8% when compared to sham controls (n=6, p<0.05; **Figure 5b**), while no other accelerations resulted in a significant difference relative to control. Taken together these data suggest that LIV can enhance proliferation in CHO-suspension cells, but that signal parameters are different than those that increase proliferation in CHO-adherent cells.

### Multiple Bouts per Day Further Enhances Proliferation of CHO Suspension Cells

Using CHO-adherent cells, we have previously reported that two LIV bouts separated by a rest period of at least 1h was more effective at enhancing proliferation than a single bout of equal total duration (Sen, Xie, et al., 2011). Here we combined the previous optimization of frequency and intensity with a 2 bout per day dosing schedule that was shown to improve LIV efficacy in adherent cells, to determine if CHO-suspension cells benefitted multiple bouts per day. CHO suspension cells were plated and allowed to culture for 24h, with one group exposed to LIV (0.7g, 30Hz, 2x1hr bout/day, 2hr refractory period) for 48h, after which cell counts were taken. LIV exposed cells showed a 210% ± 34.2% increase in proliferation when compared to sham-LIV controls (n=5, p<0.05; **Figure 6**). These data showed that more than a single bout of LIV each day, separated by a 2h refractory period, can enhance proliferation of CHO suspension cells, just as it had been shown to do in adherent cells.

**Fig. 6:**
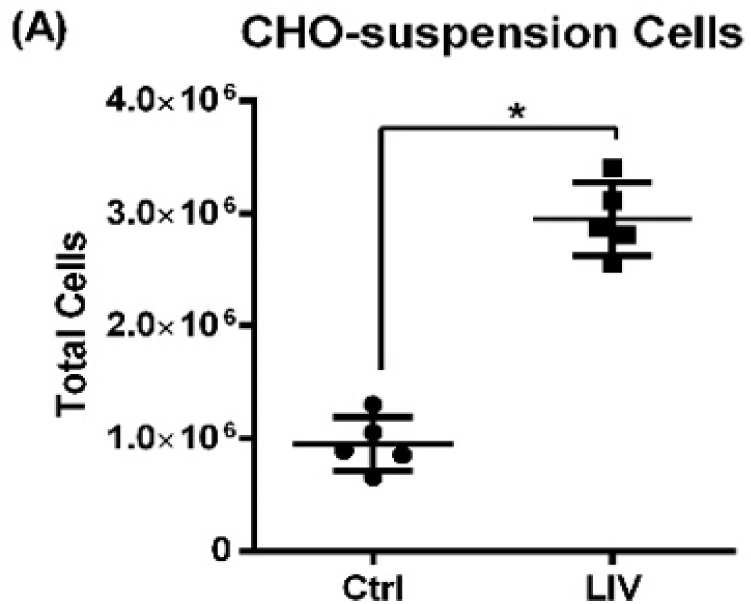
When subjected to same LIV protocol (30 Hz, 0.7g, 2 bouts of 1h/d, 2-hour refractory period), CHO-Suspension cells resulted in a 210% increase in cell proliferation. *p<0.05 Data presented as mean ± SD.

### LIV Optimization for T Cells

To determine if other types of suspension cells would respond to similar LIV parameters, a series of experiments were designed to examine if LIV could enhance T cell proliferation. Like the first optimization experiments done in CHO-suspension cells, 4 groups of T cells were subject to LIV of varying frequencies, including 0Hz (sham-handled control), 10Hz, 30Hz, and 90Hz, all at an intensity of 0.7g. After 48h of LIV, cells exposed to 30Hz signal had the greatest increase in proliferation of 24.8% ± 7.8% relative to controls (n=5, p<0.05; **Figure 7a**). Next, examining signal intensity, 4 groups of T cells were subject to LIV to include 0g (control), 0.1g, 0.7g, and 1.2g, each delivered at the ‘optimized’ frequency of 30Hz. Following 48h of treatment with the LIV protocol, cells exposed to a 0.7g acceleration had the most significant proliferative response with 20.3% ± 7.6% greater than controls. (n=6, p<0.02). Finally, examining bouts per day, experiments were designed to determine if repeated dosing with ‘optimized’ LIV frequency and acceleration resulted in an effect on T cell proliferation. T cells were separated into 4 groups: controls, 1 bout of LIV/day, 2 bouts/day, and 3 bouts/per day, with each bout separated by a 2h rest period. We observed that both 2 and 3 bouts of LIV enhanced proliferation significantly, with the 2 bouts enhancing proliferation by 22.7% ± 14.4% as compared to controls, and three bouts per day enhancing proliferation by 29.4% ± 4.3% as compared to controls (n=6, p<0.05; **Figure 7b**). Though the 3 bouts per day condition enhanced proliferation the most over control, there was no significant difference between the 2 and 3 bout conditions. Ultimately, these data suggest that LIV signal parameters, including frequency, amplitude, and number of bouts per day, can be tailored to drive a significant increase in T cell proliferation.

**Fig. 7:**
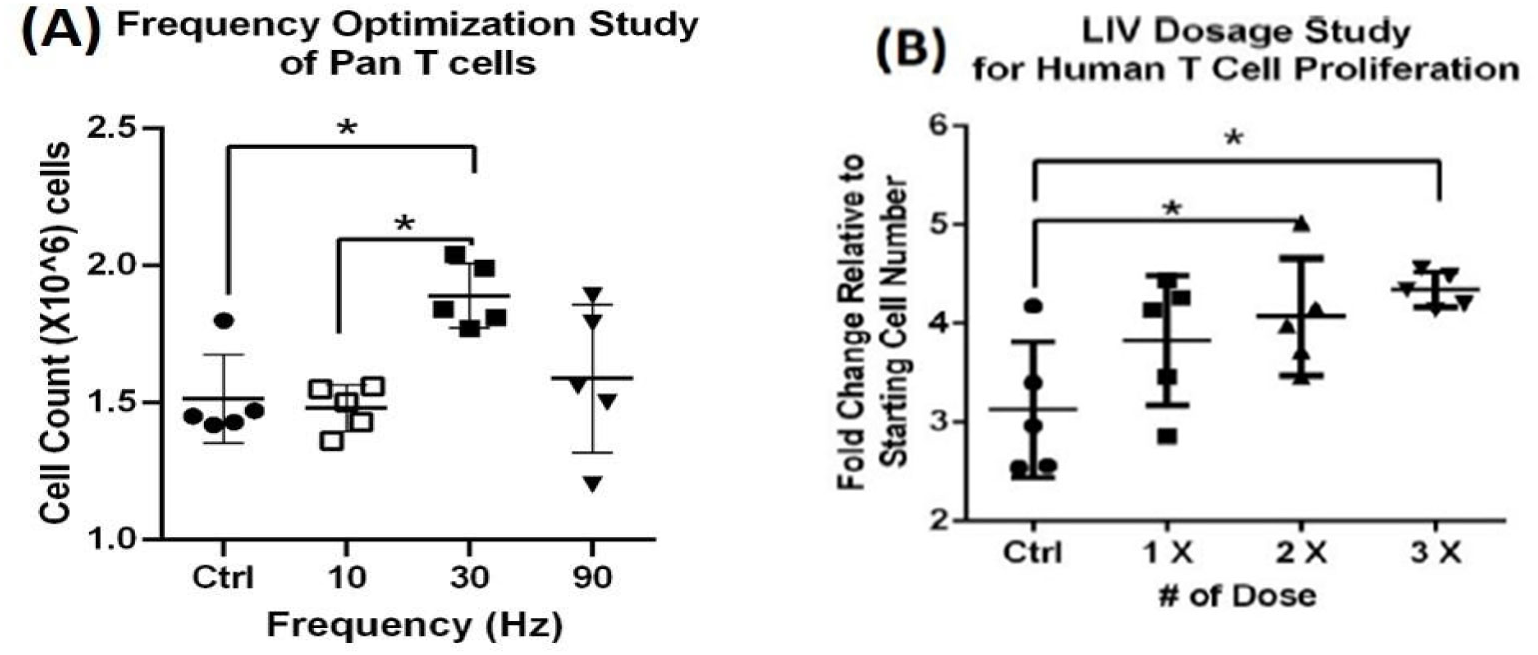
48-hour frequency and dosage optimization experiment (A) At 30Hz LIV cells showed a proliferation increase of 24.8% as compared to sham controls (n=5, mean, P<0.05). (B) Increasing from 1 to 2 LIV doses elevated proliferation by 31% (p<0.05), while 3 doses increased proliferation by 39% over sham (p=0.01)

## Discussion

Mechanical stimulation represents a relatively underutilized strategy for enhancing biomanufacturing outcomes. Our studies show that physical signals, delivered in the form of Low Intensity Vibration (LIV), can augment proliferation in both adherent and suspension cell types, but that the LIV signal is cell-type specific, with each responding differently to distinct signal configurations. That the LIV signal can be tailored to enhance expansion of both adherent and suspension cells is encouraging and suggests that LIV could be used to improve many different existing biomanufacturing processes once the influential parameters are identified.

LIV was initially developed to mimic the power spectrum of muscle contractility during exercise, as a surrogate for functional load bearing.(Huang et al., 1999) There is a wealth of literature describing the beneficial effects of exercise on a number of adherent and suspension cells within the body (Pagnotti et al., 2019), including immune cells (Yoon et al., 2021). Previous work from our lab has demonstrated that LIV can markedly influence lineage selection in MSCs *in vivo* (Rubin et al., 2007), an effect that can be translated to *in vitro* applications (Sen, Xie, et al., 2011). What was not clear was if cells in suspension – insulated from substrate strain – were also sensitive to the accelerations/decelerations of LIV, and if so, what the signal parameters would be that would drive the response. By their very definition, adherent cells are anchored to substrates via several adhesion molecules that are associated with the cytoskeleton. Since suspension cells lack this tethering, they likely perceive physical perturbations such as matrix strains and fluid shearing in a different way than adherent cells. A principal finding of the work presented here is that LIV is efficiently transmitted (>90% transmissibility) across fluids to cells in suspension, whether 24-well, 6-well plates or fully-filled T75 flasks. The only exception to this finding was the partially-filled T75 flasks where fluid movement added significantly to the LIV motions caused by the vertically oscillating elastic flask-bottom (Cox et al., 2005; Dareing et al., 2007). Therefore no experiments were conducted with partially-filled T75 flasks. When conducting experiments with CHO cells, we opted to use Erlenmeyer flasks with a substantially well supported and stiff bottom to limit wall deformations. The fact that both CHO cells and T-cells favored 30Hz response suggests that and that acceleration itself as the driving factor. These findings further indicate that if structural response of the culturing vessel is well controlled, consistently transmitting LIV to adherent and suspended cells in a scalable way is possible.

While the data reported here demonstrate that both adherent and suspended cells can respond to LIV, the outcome is highly dependent on the signal configuration, including intensity (i.e., magnitude of acceleration), frequency, bout duration, number of bouts per day and the length and number of refractory period(s). Importantly, a LIV signal tuned to stimulate expansion in one cell type is distinct from those parameters which drive another cell’s response, and thus a bioreactor using LIV to promote expansion would first have to determine the idealized signal parameters to promote a response for that given cell. Further, that suspension cells respond to LIV at all suggests that sensitivity to mechanical signals is not caused by a feature exclusive to adherent cells, or that the way they perceive mechanical signals is distinct from each other. Cells are as much an accelerometer as they are a strain gage.

One possible explanation for why suspension cells require signal parameters distinct from adherent cells could be a difference in the transmissibility of LIV. Undoubtedly, the way distortion/distention of the bottom of a flask is perceived by an adherent cell is different than how acceleration is perceived by a suspension cell. Indeed, suspension cells required a signal with a much higher amplitude (0.7g as compared to 0.2g for adherent cells) which suggests that the initial signal needs to be higher in magnitude for the suspension cells to “perceive” the same acceleration as the adherent cells. Importantly, a mechanical signal driven at a lower frequency will maintain its amplitude over longer distances and through less transmissible substrates. A signal with a higher frequency – and thus a much smaller displacement for a given acceleration - will attenuate much more quickly than a lower frequency signal.

Differences in stiffness of adherent vs suspension cell might account for their distinct responses to LIV.(Maloney et al., 2010) Given that the cytoskeleton contributes greatly to the biomechanical properties (*e.g.,* stiffness) of a cell (Discher et al., 2005), it is plausible that a “softer” cell may respond differently to LIV than a “stiffer” cell. AFM studies have provided direct evidence that mechanical connections between extracellular matrix proteins and the actin cytoskeleton indeed exist (Wang et al., 2021). The differential stiffness of adherent and suspension cells is, in part, due to their physical attachment, or lack thereof, to a surface. In attached cells, stiffness increases with development of actin fibers within a cell. Additionally, more focal adhesion complex (FAC) clusters are formed in cells attached to 3D suspended structure compared to cells attached to a 2D flat surface. The enhanced formation of focal adhesions may account for the enhanced susceptibility of adherent cells to LIV. Less efficient transmission of vibration to the nucleus might also help explain why a higher intensity was required for suspension cells to “feel” the same vibration, and suggests that conditions that disrupt the cell cytoskeleton, such as aging, will compromise mechanosensitivity.

Interestingly, previous data has confirmed that in adherent cells, a single bout of LIV will transiently induce cell-wide cytoskeletal remodeling and enhanced formation of focal adhesion clusters (Sen, Guilluy, et al., 2011). This cytoskeletal remodeling essentially “primes” a cell by improving coupling of the nucleus to the plasma edge of the cytoskeleton, and increases overall sensitivity to subsequent mechanical signals. That suspension cells also benefitted from repeated dosing with LIV, suggests that mechano-sensation in suspension cells might involve a similar type of cytoskeletal remodeling shown to play a role in adherent cells, and with enhanced cytoskeletal architecture, a ratcheted-up response to follow-on mechanical input.

Importantly we also found that LIV can enhance T cell proliferation, another type of suspension cell. It is well established that the mechanical environment plays a critical role in T cell activation and proliferation (Hu & Butte, 2016). The TCR has been classified as a mechanoreceptor and transduction of mechanical signals from the TCR is involved in modulating T cell recognition, signaling, metabolism, and gene expression (Rossy et al., 2018). Upon binding to an antigen and formation of an immunological synapse T cells also undergo dynamic reconfiguration of their cytoskeleton to better allow for precise mechanotransduction. The results reported here indicate that T cells are directly responsive to LIV and may represent a unique method for promoting T cell expansion. Additionally, it is of note that two very different types of suspension cells (CHO-suspension cells and T cells) were independently found to respond to the same set of signal parameters. This gives some support to the notion that the alteration of signal parameters between adherent and suspension cells was due to changes in transmission of the acceleration signal (i.e. altering signal parameters to generate the same changes in acceleration at the cell nucleus), and not a fundamental difference between adherent and suspension cell cytoskeletal architecture or inherent mechanosensitivity.

It is important to note that for all experiments reported here that cell proliferation was estimated by reporting daily cell counts. A more accurate analysis of changes in proliferation versus apoptosis via trypan exclusion may be warranted in addition to assessment of daily cell number. Further studies will also need to be done to determine to what effects LIV has on cell viability and functionality if this strategy should be considered for more complex biomanufacturing workflows like autologous cell therapy.

In summary, we have shown that LIV can significantly enhance proliferation of both adherent and suspended CHO cells, *in vitro,* which may translate to both biomanufacturing and autologous cell therapy industries. Our results have demonstrated that the LIV configuration is cell-type specific, meaning that what promotes proliferation in one cell type can hamper it in another. Importantly though, we have demonstrated that the LIV signal is readily ‘tunable,’ and can enhance proliferation in both adherent and suspension cells. Matrix strain and fluid shear are undoubtedly large parts of the mechanical milieu encompassing adherent cells. However, these results emphasize that suspension cells can benefit from LIV stimulation in the absence of cell anchorage, and suggests that acceleration, and not some property specific to matrix distortion of the cell, is the key driver the LIV’s effect on cell proliferation. And certainly, while the biotechnology industry has advanced bioreactors using chemical and biological approaches, there remains room for improvement. The high transmissibility of the LIV signal suggests that it can readily – and non-invasively - be incorporated into existing bioreactor infrastructure, without requiring exposing culture systems to outside agents.

## Acknowledgements

This work was supported by the Long Island Bioscience Hub NIH-Research Evaluation and Commercialization Hub (U-HL127522), the Research Foundation Technology Accelerator Fund, the Center for Biotechnology (NYSTAR) as well as grants from NIH (AG059923, P20GM109095) and NSF (1929188 & 2025505).

